# Longitudinal characterization of circulating extracellular vesicles and small RNA during simian immunodeficiency virus infection and antiretroviral therapy

**DOI:** 10.1101/2022.09.04.506571

**Authors:** Yiyao Huang, Zhaohao Liao, Phuong Dang, Suzanne Queen, Celina Monteiro Abreu, Lei Zheng, Kenneth W. Witwer

## Abstract

**Objectives:** Latent infection by human immunodeficiency virus (HIV) hinders viral eradication despite effective antiretroviral treatment (ART), Amongst proposed contributors to viral latency are cellular small RNAs that have also been proposed to shuttle between cells in extracellular vesicles (EVs). Thus, we profiled EV small RNAs during different infection phases to understand the potential relationship between these EV-associated small RNAs and viral infection.

**Design:** A well characterized simian immunodeficiency virus (SIV)/macaque model of HIV was used to profile EV-enriched blood plasma fractions harvested during pre-infection, acute infection, latent infection/ART treatment, and rebound after ART interruption.

**Methods:** Measurement of EV concentration, size distribution, and morphology was complemented with qPCR array for small RNA expression, followed by individual qPCR validations. Iodixanol density gradients were used to separate EV subtypes and virions.

**Results:** Plasma EV particle counts correlated with viral load and peaked during acute infection. However, SIV gag RNA detection showed that virions did not fully explain this peak. EV microRNAs miR-181a, miR-342-3p, and miR-29a decreased with SIV infection and remained downregulated in latency. Interestingly, small nuclear RNA U6 had a tight association with viral load peak.

**Conclusions:** This study is the first to monitor how EV concentration and EV small RNA expression change dynamically in acute viral infection, latency, and rebound in a carefully controlled animal model. These changes may also reveal regulatory roles in retroviral infection and latency.

## INTRODUCTION

Although human immunodeficiency virus (HIV) can be well controlled virologically with antiretroviral treatment (ART), chronic inflammation and other mechanisms lead to early aging disorders and serious non-AIDS events (SNAEs) including the HIV-associated neurocognitive disorders (HAND)^1,2^. Eradication of HIV reservoirs also cannot yet be achieved routinely ^3,4^. New biomarkers of SNAEs, inflammation, and responses to treatments and eradication approaches would be useful. Animal models of HIV facilitate understanding of pathology and biomarker discovery. These models allow control of ART dosing, compliance, disease monitoring, and other factors and also access to normally inaccessible compartments like the brain. Pig-tailed macaques (*Macaca nemestrina*) and rhesus macaques (*Macaca mulatta*) are well characterized SIV models of HIV disease, with some species-specific differences^5–7^.

Extracellular vesicles (EVs) are nanosized membranous vesicles released from most cells. EVs have roles in viral pathogenesis^8–10^ and, much like retroviruses, shuttle molecules between cells, including microRNAs (miRNAs)^11,12^ and small nuclear RNAs (snRNAs)^13–15^. We and others have reported correlations of miRNA expression with HIV infection, virus replication, and CNS diseases using HIV-infected CD4+ cell lines^16,17^, total plasma^18–21^, and EVs^22–26^. However, dynamic changes of EV attributes including miRNAs have not yet been fully characterized at different stages of retroviral infection and treatment.

We thus characterized EVs, EV miRNA, and U6 snRNA in longitudinal samples from pig-tailed and rhesus macaque models^27,28,29,30^, comparing pre-infection with, acute (viremic peak), latent (ART-suppressed), and rebound (ART-treatment interruption (ATI)) phases. We report that average particle counts correlate with infection, while levels of several EV miRNAs are altered even during latent infection. Remarkably, U6 snRNA is highly upregulated during infection, even in EVs purified by density gradient.

## METHODS

### Sample collection

All samples were from archives of studies approved by the Johns Hopkins University Institutional Animal Care and Use Committee and conducted in accordance with the Weatherall Report, the Guide for the Care and Use of Laboratory Animals, and the USDA Animal Welfare Act. For initial studies and verification of EV separations, plasma samples were obtained from pigtailed macaques that were not infected (n=2) or dual-inoculated with SIV swarm B670 and clone SIV/17E-Fr and untreated (n=3) or treated then treatment interruption (“rebound,” n=3); Supplemental Table 1). Longitudinal verification samples were from two cohorts of six pigtailed macaques dual-inoculated as above^27,28^ and treated with ART (consisting of once daily subcutaneous 2.5 mg/kg dolutegravir, 20 mg/kg tenofovir and 30 mg/kg emtricitabine) boosted or not with maraviroc, and a cohort of six rhesus macaques infected with SIVmac251 ^29,30^ and treated with ART (Supplemental Table 2). For pigtails, ART started at 12 days post-inoculation (dpi), and for rhesus, at 14 dpi. Time points were: pre-infection (two draws), acute infection (7, 14 dpi), latent infection (ART-suppressed) (86, 154 dpi), rebound (12 days ART treatment interruption (post-release (dpr)), and necropsy. Additional pigtailed samples (uninfected, n=3 and acute infected 7 dpi, n=3) were used for density separations (Supplemental Table 3).

### Separation of plasma EV-enriched and protein-enriched fractions

Plasma was thawed on ice and centrifuged twice at 2,500 × g (15 min, 4°C) to deplete residual platelets and debris^31,32^. 0.1 (Samples listed in Supplemental Table 2), 0.5 (Supplemental Table 1), or 8 (Supplemental Table 3) ml of platelet-depleted plasma (PDP) was separated by size-exclusion chromatography (SEC) with qEVsingle/70nm, qEVoriginal/70 nm, or qEV10/70 nm columns (Izon Science). PBS was used to elute fractions of 0.2 ml (qEVsingle/70nm), 0.5 ml (qEVoriginal/70 nm), or 5 ml (qEV10/70 nm). EV-enriched fractions (F6-8, qEVsingle/70nm; F7-9, qEVoriginal/70 nm; and F1-4, qEV10/70 nm) were pooled and concentrated by 100 kilodalton (kDa) MWCO concentrators (Thermo Fisher 88503, 88524, 88532). Pooled fractions 10-12 and 13-15 from qEVoriginal columns were collected and concentrated as protein-enriched fractions. All fractions were stored at −80°C.

### Iodixanol gradient separation

Iodixanol gradients (Optiprep™, Sigma-Aldrich D1556) were made by layering 2.5 ml each of 18%, 14%, 10%, 6% iodixanol (from 60% iodixanol at 1.320 ± 0.001 g/mL) diluted in PBS. EV-enriched fractions in 1.6 ml PBS were loaded onto the gradient. After ultracentrifugation at 200,000 × g (60 min, 4°C, TH-641 rotor, 13.2 ml thinwall polypropylene tubes, acceleration/deceleration 9), 12 fractions (0.96 ml/each) were collected from the top. Fraction densities were measured by absorbance at 340 nm. Fractions were diluted in 5 ml PBS and washed by ultracentrifugation at 200,000 × g (60 min, 4°C, AH-650 rotor, 5 ml Beckman Ultra-Clear tubes, acceleration/deceleration 9). Pellets were pipetted up and down 10 times and vortexed for 5 seconds in 120 μl PBS. Tubes were placed on ice (20 min) followed by another round of pipetting/vortexing as above. Resuspensions were stored at −80°C.

### Nanoflow cytometry (NFCM)

Concentration and size of EV- and protein-enriched fractions were measured for one minute by side-scatter using NFCM (NanoFCM) calibrated for concentration and size with 200 nm polystyrene beads and silica nanospheres, respectively (NanoFCM).

### Transmission electron microscopy

EV preparations (10 μL) were adsorbed to glow-discharged 400 mesh ultra-thin carbon-coated grids (EMS CF400-CU-UL) for 2 min followed by 3 rinses in TBS and staining in 1% uranyl acetate with 0.05 Tylose. After aspiration and drying, grids were observed with a Philips CM120 instrument at 80 kV. Images were captured by XR80 charge-coupled device (8 megapixel; AMT Imaging, Woburn, MA, USA).

### Western blotting

EV-containing fractions were lysed in 1X RIPA. Protein concentrations were determined by microBCA protein assay (Thermo Fisher, 23235). Equivalent protein amounts (EVs and proteins) were separated on 4-15% stain-free pre-cast SDS-PAGE gradient gels (Bio-Rad) under non-reducing conditions and transferred onto PVDF membranes (Sigma Aldrich). After 1 h blocking (5% non-fat milk, Bio-Rad 170–6404) at room temperature (RT), membranes were incubated with antibodies against CD63 (1:1000, BD Biosciences 556019), CD81 (1:500, Santa Cruz Biotechnology sc23962), calnexin (1:2000, Abcam ab22595), GM130 (1:1000, Abcam, ab76154), albumin (1:1000, Abcam ab28405), AGO2 (1:500, Sigma-Aldrich SAB4200085), ApoB100 (1:1000, Academy Bio-Medical 20A-G1b), ApoA1 (1:1000, Academy Bio-Medical 11A-G2b), and ApoC1 (1:1000, Academy Bio-Medical 31A-G1b) overnight at 4°C. Membranes were washed 3 times for 8 min in PBST with shaking, then incubated with HRP-conjugated secondary mouse anti-rabbit IgG or mouse IgG kappa binding protein antibodies (1:10,000, Santa Cruz Biotechnology sc-2357 and sc-516102) at RT for 1 h. After a PBST wash, membranes were incubated with SuperSignal West Pico PLUS chemiluminescent substrate (Thermo Fisher 34580) and visualized by iBright (Thermo Fisher, Waltham, MA).

### Total RNA extraction

RNA was extracted by miRNeasy Serum/Plasma Kit (Qiagen 217184) per manufacturer’s instructions, with 4 × 10^8^ copies of cel-miR-39 miRNA mimic (Qiagen 339390) spiked into to the sample after addition of lysis buffer.

### SIV Gag RNA quantification by RT-qPCR/ddPCR

Viral RNA was measured by quantitative reverse transcription-PCR (qPCR) or digital droplet PCR (ddPCR) as described ^33,34^. RNA from 140□μl of plasma was isolated by QIAamp Viral RNA Minikit (Qiagen 1020953). qPCR of SIV gag RNA was by QuantiTect Virus kit (Qiagen 211011) or ddPCR using One-Step RT ddPCR Adv kit (Bio-Rad 1864022). Copy numbers were calculated with a regression curve from control transcript standards and normalization to the volume of extracted plasma. Primers/probes for SIV gag RNA were: SIV_21_ forward 5’- GTCTGCGTCATCTGGTGCATTC-3’; SIV_22_ reverse 5’’-CACTAGGTGTCTCTGCACTATCTGTTTTG-3’; SIV_23_, 5’ FAM/3’-Black hole-labeled probe 5’-CTTCCTCAGTGTGTTTCACTTTCTCTTCTG-3 (Integrated DNA Technologies).

### SIV p27 ELISA

SIV p27 Gag was quantified by ELISA (ZeptoMetrix 0801169) per manufacturer’s instructions.

### miRNA profiling by custom TaqMan OpenArray Panel

A custom 112-assay TaqMan™ OpenArray™ MicroRNA panel (Thermo Fisher) was designed with miRNA assays chosen based on previous investigations of infectious and inflammatory diseases and identity of human and *Macaca mulatta* (mml-) miRNAs. Stem-loop primer reverse transcription and preamplification were done with manufacturer’s reagents as described^18^, but with 16 cycles of preamplification. qPCR was performed by QuantStudio 12K. Data were collected using SDS software, and quantification cycle (Cq) values were extracted with EXPRESSION SUITE v1.0.4 (Thermo Fisher Scientific). miRNAs with amplification score>1.1 and detected in > 90% of samples were included. Cq values were normalized by quantiles.

### Individual qPCR assays

Individual qPCR assays (Thermo Fisher) were performed as described^18^ for U6 snRNA (Assay ID 001973) and miRs-181a (000480), 342-3p (002260), 29a (0002112), 16 (000391), 192-5p (000491), 193b (002367), and let-7b (002619). Cq values were adjusted to the mean Cq of cel-miR-39 spike-in.

### Statistical analysis

Statistical significance of differences in EV/particle concentration, particle/protein ratio, and miRNA level between two groups were assessed by two-tailed Student’s t-test for overall ranking. Correlations between average EV/particle count and viral RNA were evaluated by Pearson’s correlation coefficient (r). Receiver operating characteristic (ROC) analyses were done in SPSS Statistics (bi-negative exponential model).

## RESULTS

### Plasma SEC fractionation

Macaque plasma was separated by SEC into early, middle, and late pooled fractions (F7-9, F10-12, and F13-15) and characterized per MISEV recommendations^31^. Despite similar particle concentrations (Figure S1A), F7-9 had significantly less protein versus other fractions, consistent with relatively pure EVs. Transmission electron microscopy (TEM) showed cup-shaped oval/round particles in F7-9 but non-EV aggregates in others (Figure S1B). Western blot for EV-enriched membrane markers CD63 and CD8; argonaute protein 2 (AGO2, which may be present at low levels in EVs but is mostly outside EVs in plasma); cellular markers GM130 and calnexin; and albumin and lipoproteins ApoB100, ApoA1, and ApoC1 (Figure S1C and D). We thus defined F7-9 as EVs, and F10-12 and F13-15 as proteins.

### Particle counts and sizes in uninfected, infected, and rebound groups

EVs and proteins were characterized from a small number of uninfected (n=2), infected and untreated (44-49 dpi, n=3), and rebound (infected/ART treatment interrupted) (Supplemental Table 1) subjects. Five of six infected samples from untreated and rebound groups had greater particle counts than un-infected in EV (though not statistically different due to the small sample size used) but not protein fractions (Figure 1A). The EV fractions of infected, untreated animals had smaller average particle sizes, but following treatment interruption, there were more EVs with diameter >120 nm (Figure 1B). Since the majority of particles had diameters < 100 nm, intact virions (100-130 nm) could not fully explain these differences.

**Figure 1.**
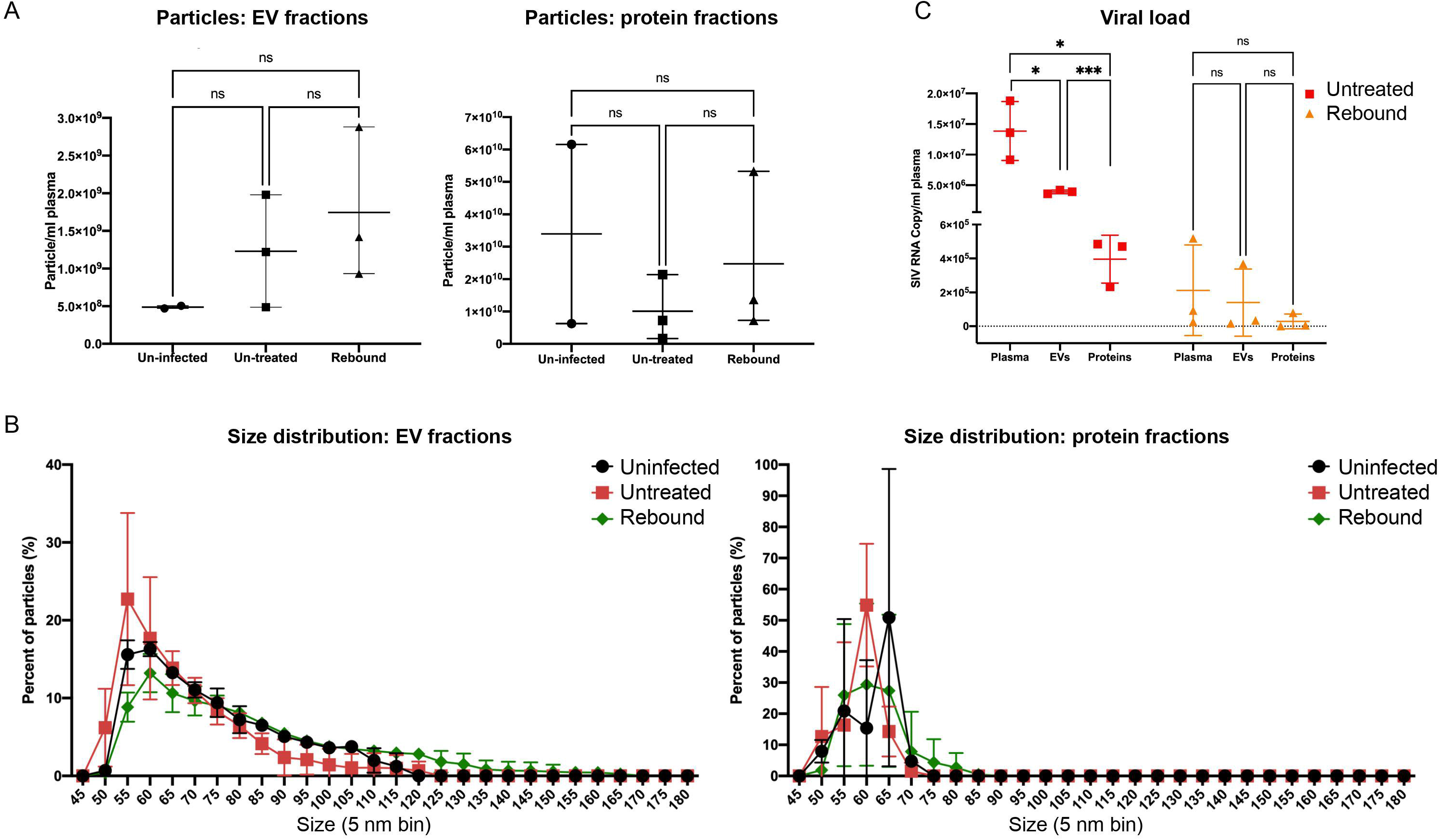
Characterization of plasma EV- and protein-enriched fractions from SIV-uninfected and infected non-human primate models. (A) Particle concentrations of EV- (left) and protein- (right) enriched fractions of uninfected, untreated, and rebound animals were measured by HSFCM. Particle concentration for each group was normalized by plasma input (per 1 ml). (B) Size distributions of EV (left) and protein (right) fractions were measured by HSFCM and calculated as particles in a specific size bin versus total detected particles in each sample (percentage). (C) Viral load (GAG RNA qPCR) as copy number per ml plasma for unfractionated plasma, EVs, and proteins. Data are mean +/− SD. ns: no significant difference (p > 0.05), *p ≤ 0.05, ***p ≤ 0.001 by two-tailed Welch’s t-test.

### Distribution of SIV gag RNA

One third of total plasma viral RNA was recovered in EV fractions (Figure 1C): one order of magnitude greater than recovery from protein fractions. During rebound, most plasma gag RNA (mean 2.1 × 10^5^ copies/ml) was recovered in EVs (mean 1.4 × 10^5^ copies/ml): two orders of magnitude greater than recovery from protein (mean 2.81 × 10^3^ copies/ml).

### Longitudinal particle and viral RNA concentrations in three HIV/SIV models

We next examined species, SIV strain, and ART regimens with three cohorts of NHP (n=6 each, Supplemental Table 2; viral load, Figure S2). Group A was pigtailed macaques receiving ART at 12 dpi. For Group B, ART was augmented with CCR5 inhibitor Maraviroc. Group C was rhesus macaques treated with ART at 14 dpi. Combining all subjects, average particle concentration in EV fractions was positively correlated with average plasma SIV RNA throughout infection phases (R=0.8916, p < 0.05; Figure 2A). However, the correlation of particle concentration and viral RNA was significant only in Group A (R=0.88, p < 0.05; Figure 2B), nearing nominal statistical significance in Group B (R=0.71, p=0.07; Figure 2B) but not Group C (R=-0.12, p=0.79; Figure 2B; see also Figure 2C for individual particle counts). Average viral loads at 7dpi were highest for Group A, followed by B and C (Figure 2B). Virions, which comprise only a small percentage of total particles, do not explain these differences. Size distribution in EV fractions was similar between groups and time points, with the majority of detected particles <100 nm in all groups (Figure S3).

**Figure 2.**
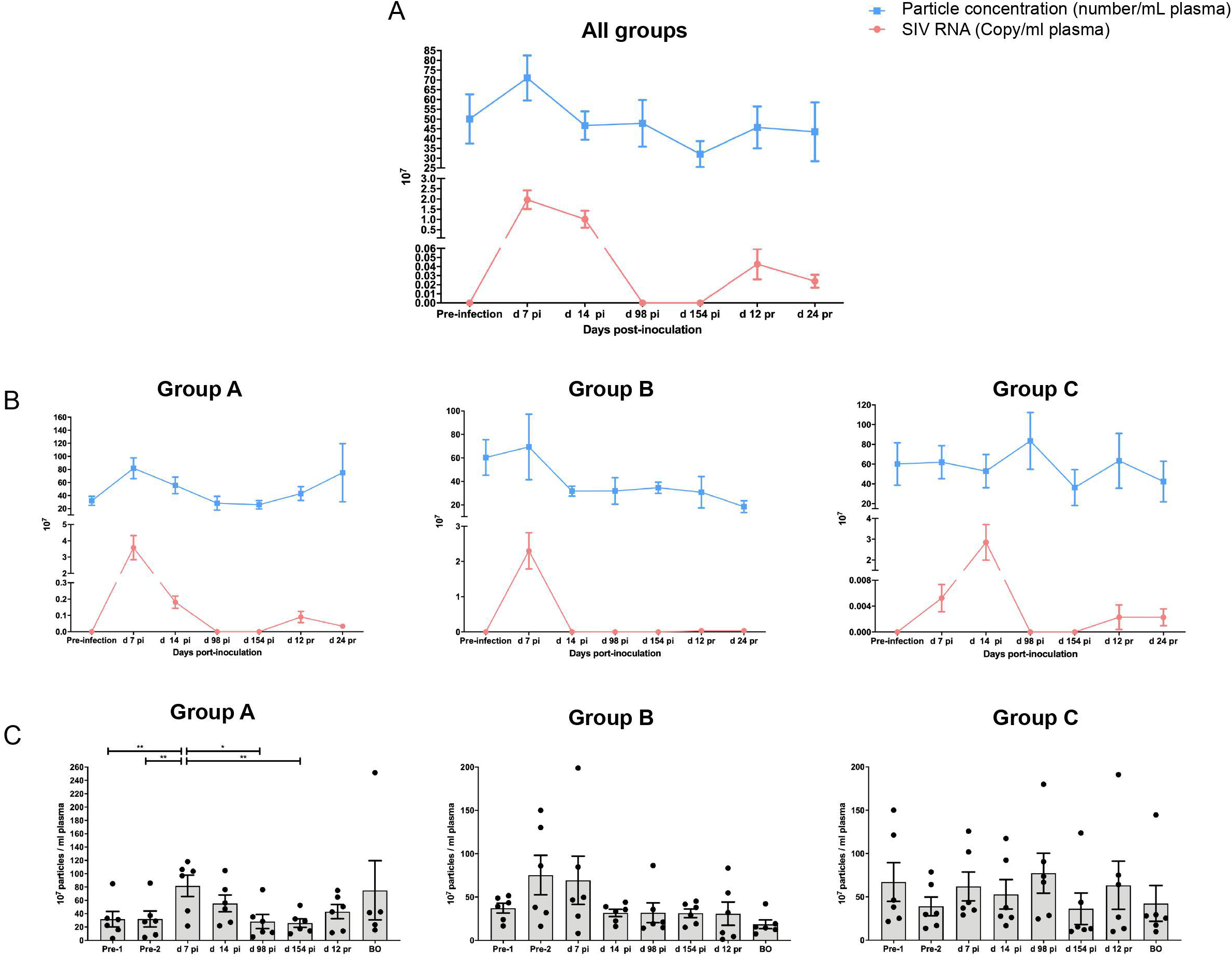
Plasma particle concentration and viral load in longitudinal samples from nonhuman primate (NHP) models. Plasma EV particle concentration measured by HSFCM and SIV GAG RNA copy number (qPCR) of eighteen NHPs (A) and NHPs separated by groups based on species and treatment regimen (B). Particle concentration and viral RNA copy number for each group was normalized by plasma input (per 1 ml). (C) Particle concentrations of EV-enriched fractions at different infection phases were measured by HSFCM. Particle concentration for each group was normalized by plasma input (per 1 ml). Data are mean +/− SD. *p ≤ 0.05, **p ≤ 0.01, ***p ≤ 0.001, ****p ≤ 0.0001 by two-tailed Welch’s t-test.

### Plasma EV sRNA profiles during SIV infection

EV-associated small RNAs (sRNAs, mostly miRNAs) were measured by custom microarray. 56 features satisfied inclusion criteria. Significantly differentially abundant sRNAs with fold change >1.5 in acute infection, latent infection (ART-suppressed), and rebound (after treatment interruption) vs pre-infection are shown in Figure S4A. Most sRNA differences were found in Group A, with the largest acute and rebound viral loads. Similarly, more sRNAs were dysregulated during acute infection than in other phases (Figure S4A). Several sRNAs were consistently dysregulated across two or more groups. During acute infection, these included miR-29a and miR-145 (Groups A,B) and miR-342-3p and U6 (all groups). During ART treatment, there were no consistent sRNA changes compared with pre-infection. During viral rebound, miR-192 (A,B) and miR-146b and miR-342-3p (A,C) were partly consistent. Thus, the only consistent changes across groups were of miR-342-3p and U6 during acute infection.

### EV miRNA dysregulation: individual qPCR validation

Individual qPCR assays confirmed downregulation of miRs-342-3p, 181a, and 29a after SIV infection in pigtails (A,B, Supplemental Table 2) (Figure 3A). miRs-342-3p and 181a were not only dysregulated in acute infection, but also remained downregulated during latent and rebound phases. To assess association with phase, receiver operating characteristic (ROC) curves were generated (Figure 3B). miR-342-3p alone had greater area-under-the-curve (AUC, 0.83±0.10) in discriminating pre-infection from rebound, while a combination of three miRNAs had greater AUC in other comparisons: 0.77±0.10 (acute 7 dpi), 0.81±0.10 (acute 14 dpi), 0.75±0.10 (latency). We also examined how these miRNAs changed dynamically in the 12 subjects individually (Figure S5A). Despite large inter-subject variation, miRs-342-3p, 181a, and 29a were downregulated after SIV infection in at least half of the subjects. In contrast, for rhesus macaques (Figure S5B, Group C), only miR-29a was differentially abundant between acute infection (14 dpi) and latent infection but largely attributed to one outliers (Figure S5C).

**Figure 3.**
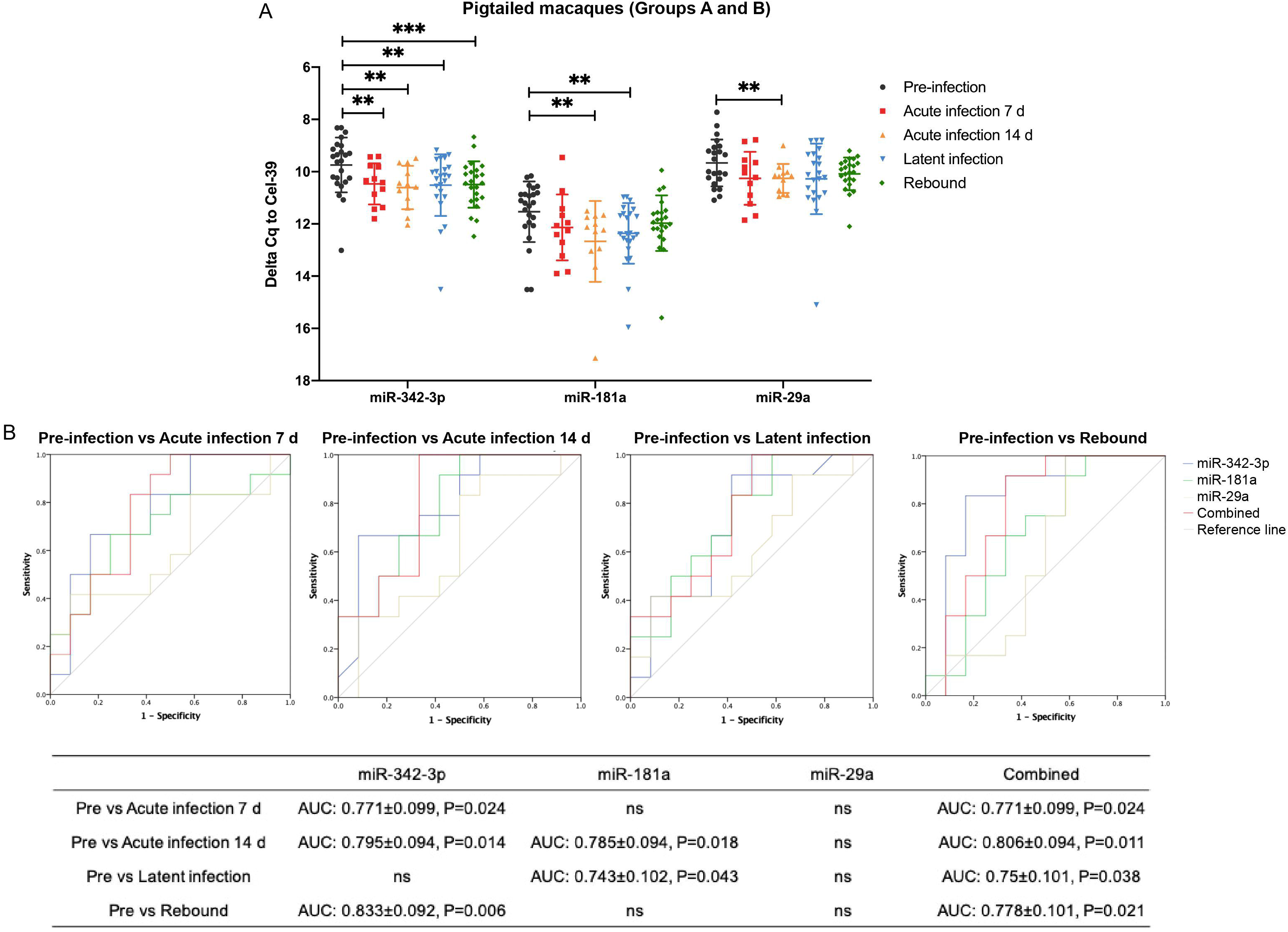
EV miRNA individual qPCR validation in pigtailed macaques. (A) qPCR validation for miR-342-3p, miR-181a, and miR-29a in pigtailed macaques (n = 12) at different infection phases. Delta Cq values was normalized to the spike in cel-miR-39 control. Data are mean +/- SD. *p ≤ 0.05, **p ≤ 0.01, ***p ≤ 0.001, ****p ≤ 0.0001 by two-tailed Welch’s t-test. (B) Receiver operating characteristic (ROC) curves for the levels of three individual EV miRNAs and a combined three-miRNA panel to differentiate pre-infection from acute infection (7 and 14 dpi), latency, and rebound.

### U6 snRNA correlates with plasma viral RNA peak in pigtailed and rhesus macaques

U6 is a commonly used reference in miRNA qPCR assays, especially for examination of cellular and tissue RNA. EV-associated U6 snRNA increased in both pigtailed and rhesus macaques during peak viral load in acute infection (7 dpi, pigtails, 14 dpi, rhesus, Figure 4A; viral load, Figure S2), with log2 fold changes versus pre-infection of 2-3 on average (Figure 4B). Tracking the abundance of EV U6 snRNA across infection (Figure 4C), changes in EV U6 levels were highly consistent. Association with infection was also supported by ROC curves (Figure 4D).

**Figure 4.**
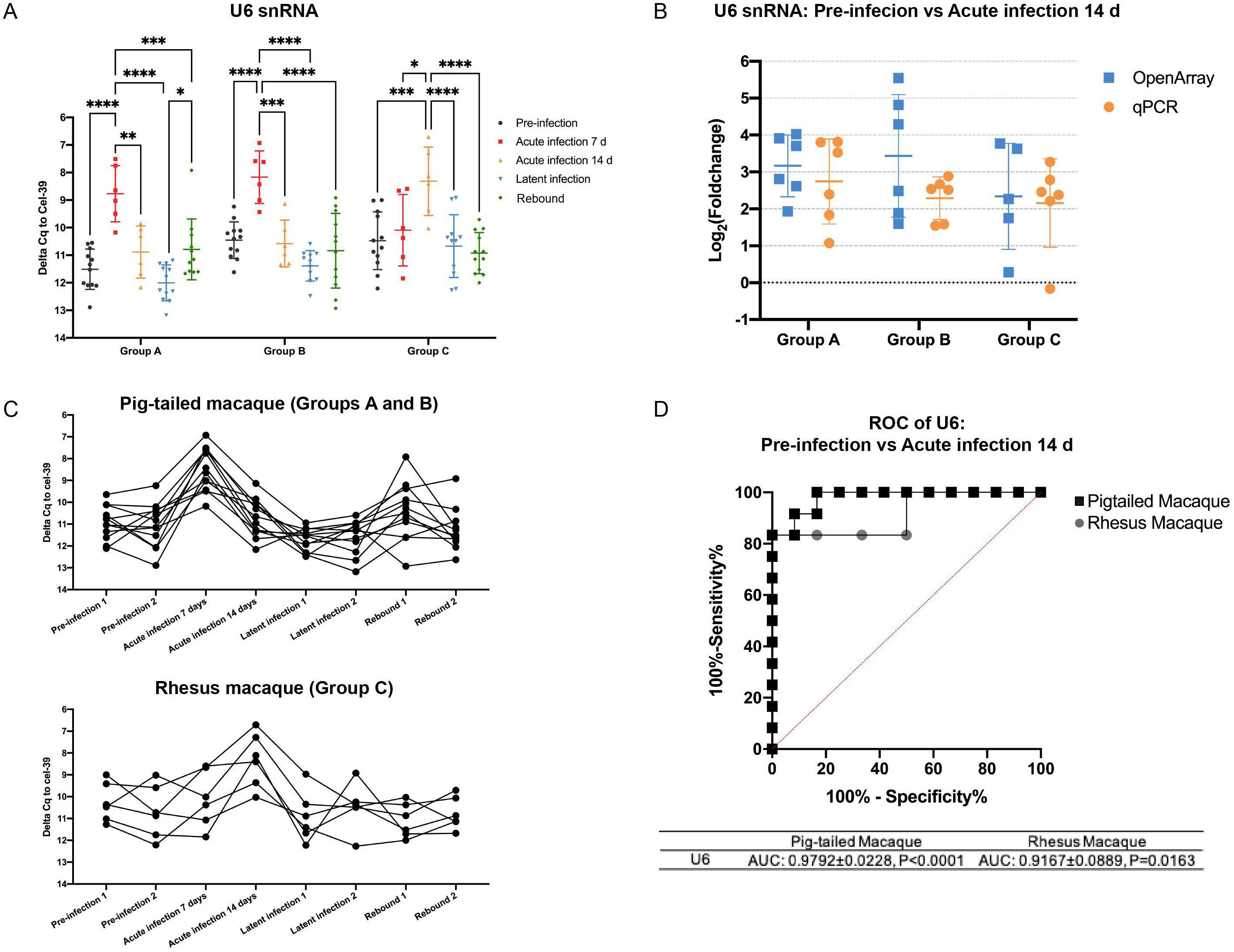
EV U6 snRNA level is tightly associated with the acute infection phase as validated by individual qPCR. (A) Individual qPCR validation of U6 snRNA in Groups A, B, and C at different infection phases. Delta Cq values were normalized to a spiked-in cel-miR-39 control. Data are presented as mean +/− SD. *p ≤ 0.05, **p ≤ 0.01, ***p ≤ 0.001, ****p ≤ 0.0001 by two-tailed Welch’s t-test. (B) Log2 (fold change) of EV U6 snRNA in acute infection compared to pre-infection from OpenArray and individual qPCR analyses. Data are mean +/− SD. (C) U6 snRNA levels in pigtailed and rhesus macaques in longitudinal samples. Delta Cq values were normalized to cel-miR-39 control. (D) Receiver operating characteristic (ROC) curves for the levels of EV U6 to differentiate pre-infection from acute infection in pigtailed and rhesus macaques.

### U6 snRNA in gradient-separated EVs

Increased U6 in EV fractions during infection could be due to packaging into EVs, virions, or both. We thus further fractionated EVs from uninfected and acute infected samples (Supplemental Table 3) into lighter EV fractions and denser virus populations using differential gradient ultracentrifugation with iodixanol. For acute infected samples, SIV p27 Gag protein (Figure 5A) and RNA (Figure 5B) were determined for input plasma and EV-enriched SEC fractions. Consistent with the findings in Figure 1, not all viral protein and RNA was recovered in the EV fractions, likely because some material is lost to the column matrix. Also as before, total particles by NFCM tended to be greater in infection (Figure 5C). qPCR confirmed that miRs-342-3p and 29a were less abundant, while U6 snRNA was more abundant, in acute infection samples, although miR-181a abundance was similar (Figure 5D).

**Figure 5.**
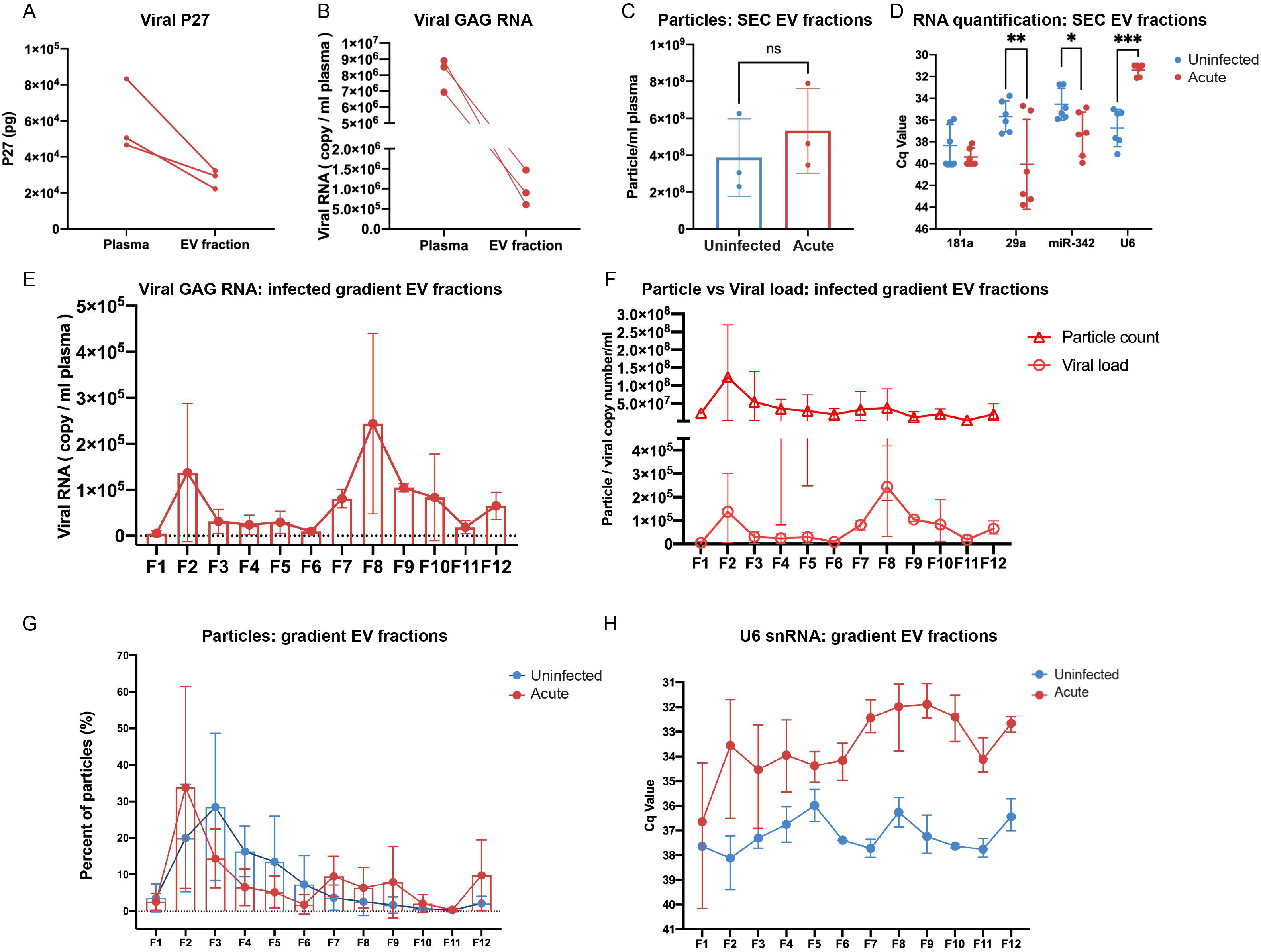
EV and EV small RNA characterization in iodixanol density fractions. SIV P27 GAG protein (A) measured by ELISA and GAG RNA (B) by qPCR in raw plasma and EV enriched SEC fraction from SIV acutely infected pigtailed macaque plasma (n=3). (C) Particle concentrations of EV-enriched SEC fractions of uninfected (n=3) and acutely infected (n=3) pigtailed macaques as measured by HSFCM. Particle concentration for each group was normalized by plasma input (per 1 ml). Data are mean +/− SD. ns: no significant difference (p > 0.05) by two-tailed t-test. (D) qPCR validation for miRNAs and U6 snRNA in EV-enriched SEC fractions from uninfected and acutely infected plasma. Data are mean +/- SD. *p ≤ 0.05, **p ≤ 0.01, ***p ≤ 0.001, ****p ≤ 0.0001 by two-tailed t-test. (E) GAG RNA level detected by qPCR in 12 EV fractions separated by iodixanol density gradient in acutely infected plasma (n=3). Data are mean +/− SD. (F) EV particle concentration and SIV GAG RNA copy number were plotted for 12 EV iodixanol fractions of acutely infected plasma (n=3). Particle concentration and viral RNA copy number for each group was normalized by plasma input (per 1 ml). Data are mean +/− SD. (G) Particle number distribution of 12 iodixanol fractions from uninfected (n=3) and acutely infected plasma (n=3). Particle concentration for each fraction was measured by HSFCM and calculated as particles in each fraction versus total particles recovered from 12 fractions (percentage). Data are mean +/− SD. (H) The level of U6 in 12 iodixanol fractions from uninfected (n=3) and acute infected plasma (n=3). Data are mean +/− SD.

Following gradient separation, the density distribution of 12 collected fractions was similar for uninfected and acute infected samples, ranging from 0.961 to 1.187 g/ml (Figure S6A). Particles with EV morphology were found by TEM in all fractions, although it was difficult to identify virions (Figure S6B). Transmembrane protein CD9 was detected in F2-9 (acute infected), and F2-4 (uninfected; Figure S6C). The pattern of CD9 distribution for the acute infected sample suggests successful separation of a light EV fraction (~F2-4) and a heavy EV and/or virion fraction (~F7-9). Gag RNA was detected in most fractions with a peak around F8, consistent with presence of virions, while Gag protein was below the limit of detection (Figure 5E and data not shown). There was no significant correlation between particle concentration and viral RNA (R=0.3501, p=0.2645; Figure 5F). Particle recovery was greatest in F2 and F7-F9 (acute infected), but in F3 for uninfected, with few particles in F7-9 (Figure 5G). U6 snRNA was more abundant in all infected fractions except for F1, with the highest average abundance in the heavier F7-10 (Figure 5H). miRs-342-3p and 29a, while detected, were highly variable (Figure S6B).

## DISCUSSION

As the gold standard to measure the latent HIV reservoir, quantitative virus outgrowth assays (qVOAs) are laborious, expensive, and require a large amount of blood samples^35^. The reservoir could potentially be measured using EVs, with contents reflecting disease status and easily accessible in blood. Here, for the first time, we examined EV concentration and small RNA contents in longitudinal samples from SIV models, finding that EV concentration, miRNA, and U6 snRNA levels are linked to SIV infection. Density separation of EV subtypes revealed strong enrichment of U6 snRNA in all EV subtypes in acutely infected versus uninfected plasma.

To be sure, overlapping contents and physical properties make EVs and virions (or “host EVs” and “viral EVs”) difficult to separate^36,37,38,39^. While density gradients are a standard method for virion/EV separation^40–44^, most methods have been optimized for cell culture medium^41–43^ or large quantities of virions spiked into plasma^44^. However, our EV and EV exRNA results are likely not skewed by the presence of virions. First, retrovirions are less abundant than EVs in biofluids, even in uncontrolled viral replication. We observed a significantly lower viral GAG RNA copy number than nanoparticle number in EV-enriched SEC fractions, even during viral peak. Both GAG RNA and GAG antigen levels were significantly lower in EV-enriched SEC fractions versus unfractioned plasma. In addition, gradient-separated EVs did not contain detectable GAG antigen but did have GAG RNAs. Second, GAG RNA was also detected in protein fractions, with particles of much smaller diameter than virions. Finally, viral RNA fragments could also be incorporated into host EVs^45,46^. Thus, increased particles in infected plasmas are not explained solely by retrovirions.

Previous studies reported increased plasma EV concentration in HIV-infected patients compared with uninfected controls^23,24,47,48^, albeit with no differences between ART-naïve and ART-suppressed patients^23^. In some cases, EVs were quantitated by acetylcholinesterase (AChE)^48^, which may not be specific for EVs^49^. Moreover, separating EVs from e.g., lipoproteins is difficult using ultracentrifugation or polymer precipitation^31^. Here, we used size exclusion chromatography (SEC), a good option for obtaining relatively pure EVs^50,51^. We also further separated EV subtypes and virions by gradient, revealing EV and EV small RNA distributions. Although we cannot rule out all co-isolates, and thus refer to “EV-enriched fractions,” our results support more EV production during SIV infection. Even so, changes appeared to vary with species and individual subject, and more work is needed to understand mechanisms behind this and previous findings on release of EVs after retroviral infection^52,53,54^.

Where do our miRNA findings fit into the existing literature? We previously reported that total plasma miRNA profiles change during acute SIV infection^18^ and that EV-borne miRNA miR-186-5p is downregulated in cervicovaginal lavage (CVL) and regulates HIV replication in macrophages^55^. Here, we identified several miRNAs that were dysregulated during latent and rebound infections, but only very few compared with the acute infection phase, consistent with a study comparing productive and latent infection in CD4+ T cell models^16^. Some identified miRNAs were previously reported in HIV infection or inflammation, such as miR-29a^56–59^ and miR-181a^16,60,61^, while miR-342-3p was not. However, unlike particle concentration, which varied with viral load, EV miRNAs remained downregulated after infection even when viral RNA was undetectable. Of note, host miRNA changes are probably not specific to HIV infection. For example, miR-29a, identified by us and other groups, was previously observed to be down-regulated in tuberculosis^62^ and hepatitis C virus (HCV)^57^ infection. EV miRNA may thus be useful to indicate persistent infection generally but may not be a good indicator for specific disease monitoring.

Our findings on U6 were unexpected and potentially informative. U6 snRNA is one of the most highly conserved RNAs across species and is thus commonly used as an internal control gene in, e.g., miRNA qPCR assays^63–66^ despite some subject- and disease-related differences^67–71^. Although a few studies revealed upregulation of EV U6 in cancer and smoking^72–74^, such changes have not to our knowledge been reported in infectious diseases. Our data suggest that U6 is not an ideal reference RNA, at least in retroviral infection. Interestingly, U6 snRNA is also one of the host cellular RNAs reported to be specifically packaged into retroviral particles^75–77^, for example, in murine leukemia virus (MLV) infection^75,78^. Retroviral packaging of U6 may be independent of viral full-length RNA transcripts^77,79^ but affected by the nucleocapsid (NC) domain of the Gag polyprotein^79^. However, standard virus purifications^76,77,79^ also co-isolate EVs. How U6 is packaged into EVs is unclear, as is the possible function of this RNA in the cell or in the EV during retroviral infection.

In summary, our rigorous EV separation and characterization strategy showed that plasma EVs and several EV sRNAs change consistently during the course of SIV infection across several related models. Our findings raise new questions about the distribution of host RNAs released in “host” EVs compared with “hijacked” EVs (retrovirions) and sound a cautionary note about the use of U6 as a reference RNA, at least in retroviral infection. We are now applying these methods to studies of human plasma to understand if EV RNAs will provide value in monitoring HIV latent infection.

## ACKNOWLEDGMENTS

The authors thank members of the Witwer Laboratory for discussions and support. We are particularly grateful to members of the Retrovirus Laboratory for access to samples from the animal models and for helpful suggestions. Electron microscopy images were acquired in the Johns Hopkins University School of Medicine Institute for Basic Biomedical Sciences Microscope Facility. The authors would like to thank ViiV Pharmaceuticals for their generous donation of dolutegravir and maraviroc, and Gilead Pharmaceuticals for their generous donations of tenofovir and emtricitabine. This work was supported in part by the US National Institutes of Health, National Institute on Drug Abuse (NIDA, DA040385 and DA047807 to KWW), by National Institute of Mental Health Grant No.P30MH075673 to the Johns Hopkins NIMH Center, and by the Johns Hopkins University Center for AIDS Research Grant No. P30AI094189 (pilot grants to YH). The Witwer lab is also supported in part by NCI/Common Fund CA241694, NIAID AI144997, NIMH MH118164 (to KWW), the Richman Family Precision Medicine Center of Excellence in Alzheimer’s Disease, U42OD013117 (Johns Hopkins, NIH-supported pigtailed macaque breeding colony), NINDS NS089482 (to Joseph L. Mankowski), and NIMH MH070306 (to Janice Ellen Clements).

---

Figure S1 Characterization of plasma EV- and protein-enriched fractions separated by size exclusion chromatography (SEC). (A) Particle (left) and protein (middle) concentrations of pools of Fractions F7-9, F10-12, and F13-15 from SEC were measured by HSFCM and microBCA protein assay. Particle and protein concentration for each group was normalized by plasma input (per 1 ml). Right: Ratio of particles to protein (particles/μg). Data are mean+/− SD. *p ≤ 0.05, **p ≤ 0.01, ***p ≤ 0.001, ****p ≤ 0.0001 by two-tailed Welch’s t-test. (B) SEC fraction pools were visualized by negative staining transmission electron microscopy (scale bar = 100 nm). TEM is representative of five to ten images taken of each fraction from three independent plasma separations. (C) Western blot of GM130, calnexin, Ago2, CD63, and CD81 associated with SEC fractions. Cell lysate from dendric cells and 12,000 × g-centrifuged plasma supernatants (12k sup) were used as positive and negative controls. (D) Western blot of ApoB100, ApoA1, ApoC1, and albumin associated with SEC fractions. Cell lysate from dendric cells and another SEC-based method (called smartSEC) were used as positive and negative controls.

Figure S2 Longitudinal viral loads of plasma samples from pigtailed and rhesus macaques. 12 pigtailed macaques were infected with SIV swarm B670 and SIV/17E-Fr, while six rhesus macaques were infected with SIVmac251.

Figure S3 Size distributions of EV-enriched fractions in groups A, B, and C collected at different infection phases were measured by HSFCM and calculated as particles in a specific size bin versus total detected particles in each sample (percentage). Data are presented as mean +/− SD.

Figure S4 EV miRNA dysregulation identified by OpenArray in acute viral infection, latent, and rebound in SIV models. (A) Significantly differential expressed miRNAs with fold change > 1.5 in acute, latent, and rebound infection as compared with pre-infection. (B) Venn diagrams of dysregulated miRNAs in acute, latent, and rebound infection as compared to pre-infection identified in the A and C groups.

Figure S5 EV miRNA qPCR validation. (A) miRNAs as measured by qPCR in longitudinal plasma of pigtailed macaques. Delta Cq values was normalized to the spiked-in cel-miR-39 control. (B) qPCR validation for EV miR-29a in rhesus macaques (n = 6) at different infection phases. Data are mean +/− SD. *p ≤ 0.05 by two-tailed Welch’s t-test. (C) The level of EV miR-29a as measured by qPCR in longitudinal plasma of rhesus macaques. Delta Cq values were normalized to the spiked-in cel-miR-39 control.

Figure S6 EV characterization of iodixanol gradient fractions. (A) Density distribution of 12 iodixanol fractions from uninfected (n=3) and acutely infected plasma (n=3). Data are mean +/− SD. (B) Iodixanol fractions visualized by TEM (scale bar = 100 nm). TEM is representative of five to ten images taken of each fraction from one pair of uninfected and infected samples. (C) Western blot of CD63 and CD9 associated with iodixanol fractions from one pair of uninfected and infected samples. (D) qPCR validation for miR-342-3p and miR-29a in iodixanol fractions from uninfected and acutely infected plasmas. Data are mean +/− SD.

