## Supplemental Figures for "Longitudinal characterization of circulating extracellular vesicles and small RNA during simian immunodeficiency virus infection and antiretroviral therapy"

Figure S1

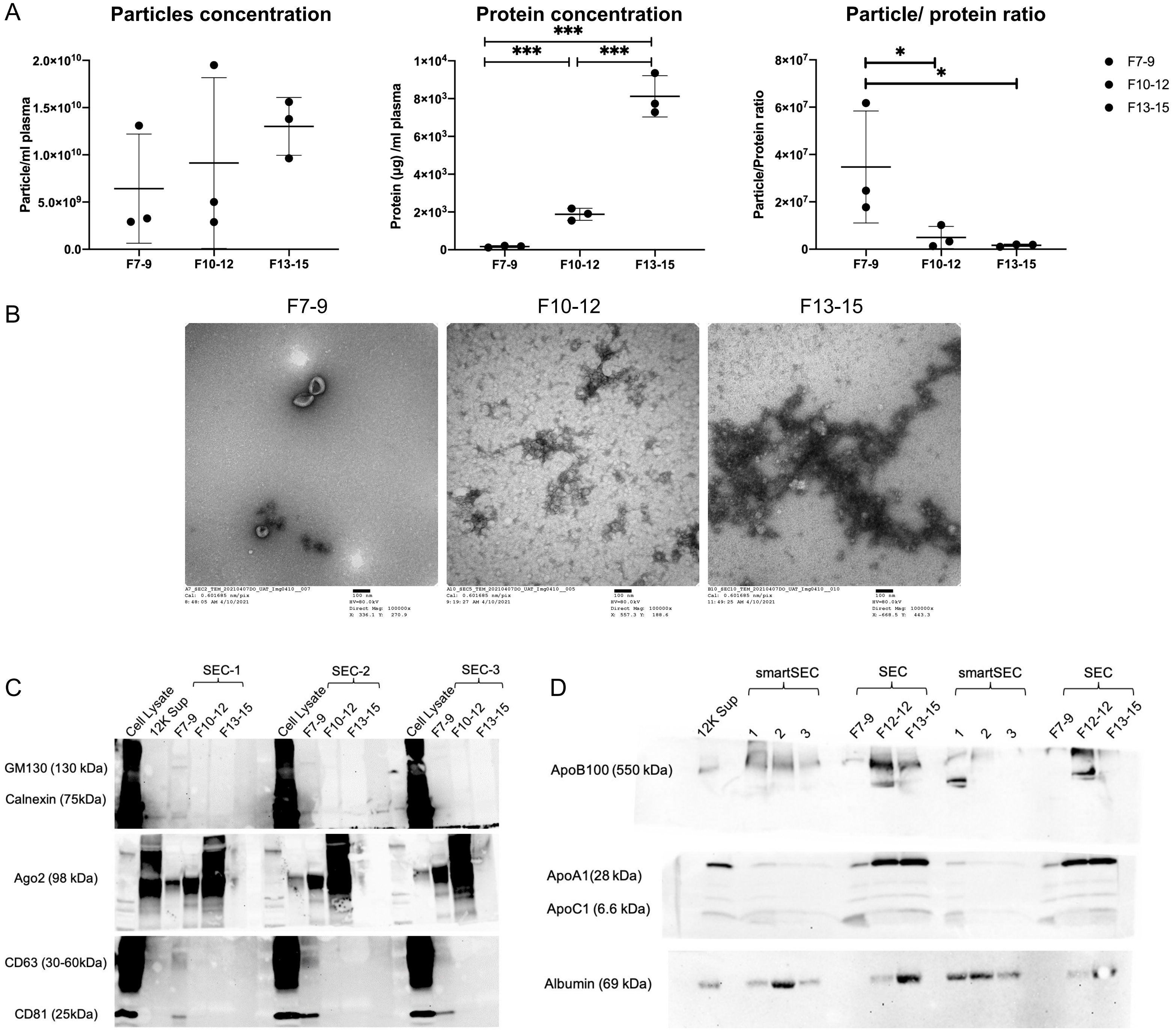

A
Particles concentration
Particle/ml plasma
2.0×10^10
1.5×10^10
1.0×10^10
5.0×10^9
0.0
F7-9
F10-12
F13-15
Protein concentration
Protein (µg) /ml plasma
1×10^4
8×10^3
6×10^3
4×10^3
2×10^3
0
***
Particle/ protein ratio
Particle/Protein ratio
8×10^7
6×10^7
4×10^7
2×10^7
*
F7-9
F10-12
F13-15
B
F7-9
F10-12
F13-15
C
SEC-1
SEC-2
SEC-3
Cell Lysate
12K Sup
GM130 (130 kDa)
Calnexin (75kDa)
Ago2 (98 kDa)
CD63 (30-60kDa)
CD81 (25kDa)
D
smartSEC
SEC
12K Sup
1
2
3
F7-9
F12-12
F13-15
ApoB100 (550 kDa)
ApoA1(28 kDa)
ApoC1 (6.6 kDa)
Albumin (69 kDa)

Figure S2

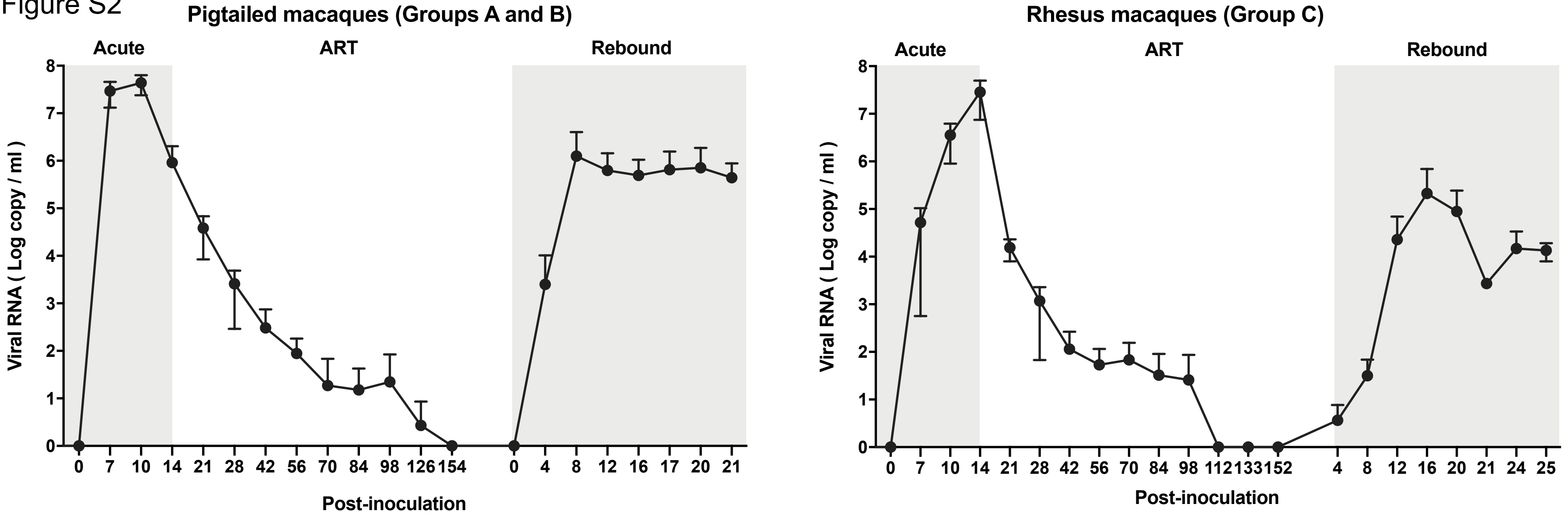
Pigtailed macaques (Groups A and B)
Acute
ART
Rebound
Viral RNA ( Log copy / ml )
0
1
2
3
4
5
6
7
8
0 7 10 14 21 28 42 56 70 84 98 126 154 0 4 8 12 16 17 20 21
Post-inoculation
Rhesus macaques (Group C)
Acute
ART
Rebound
Viral RNA ( Log copy / ml )
0
1
2
3
4
5
6
7
8
0 7 10 14 21 28 42 56 70 84 98 112 133 152 4 8 12 16 20 21 24 25
Post-inoculation

Figure S3

**Group A**

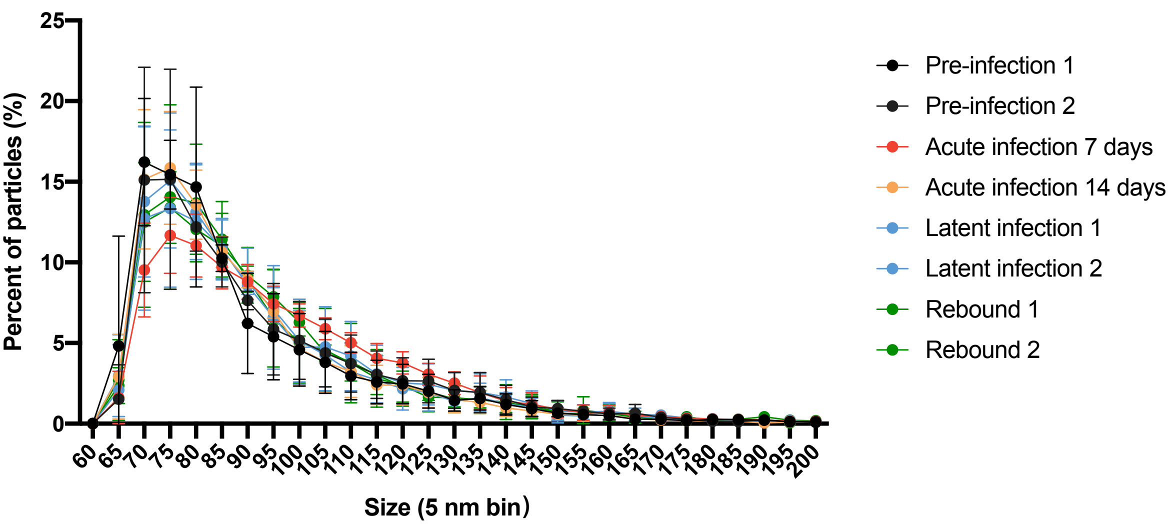

**Group B**

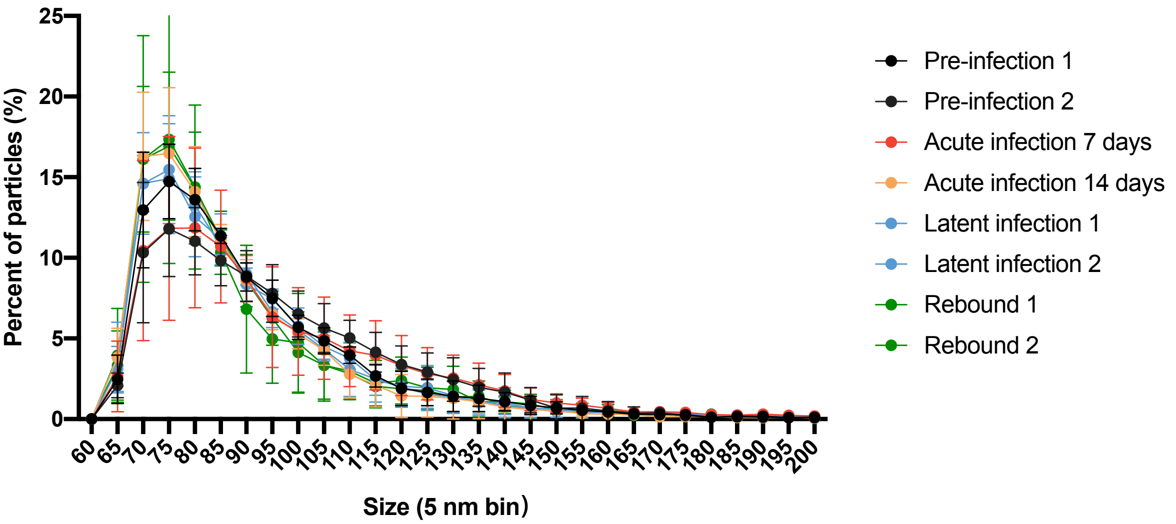

**Group C**

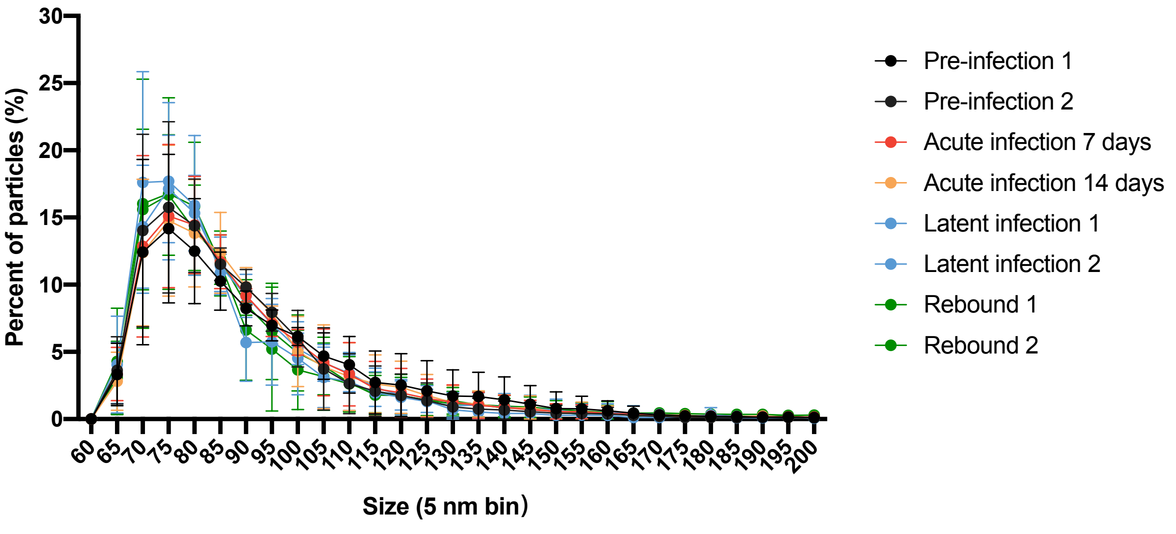

Figure S4

A

| | Group A | Group B | Group C |
|---|---|---|---|
| | miR-342-3p | miR-342-3p | miR-342-3p |
| | U6 snRNA | U6 snRNA | U6 snRNA |
| | miR-29a-3p | miR-29a-3p | |
| | miR-145-5p | miR-145-5p | |
| | miR-181a-5p | | |
| **Pre vs Acute** | miR-193b-3p | | |
| | miR-195-5p | | |
| | miR-19a-3p | | |
| | miR-328-3p | | |
| | miR-451-5p | | |
| | | | miR-638-5p |
| | miR-125b-5p | | |
| | miR-146a-5p | | |
| **Pre vs Latent** | miR-181a-5p | | |
| | miR-21-5p | | |
| | | let-7c-5p | miR-342-3p |
| | miR-192-5p | miR-192-5p | |
| | miR-342-3p | | miR-342-3p |
| | miR-146b-5p | | miR-146b-5p |
| | miR-125b-5p | | |
| **Pre vs Rebound** | miR-145-5p | | |
| | miR-181a-5p | | |
| | let-7c-5p | | |
| | | miR-454-3p | |
| | | | miR-638-5p |
| | | | miR-1290-3p |

B

Group A （n=6）

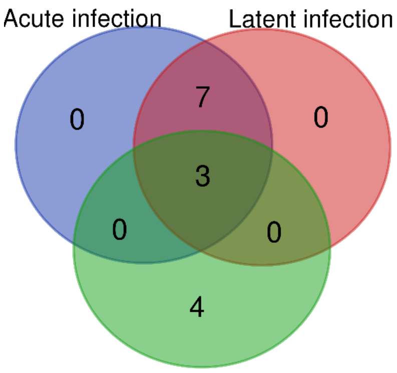

C

Group C （n=6）

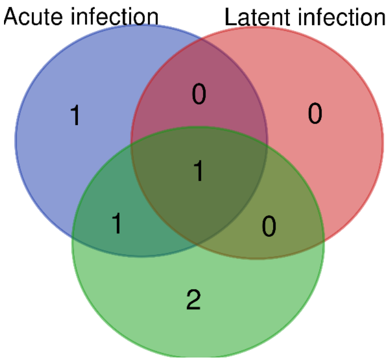

Figure S5

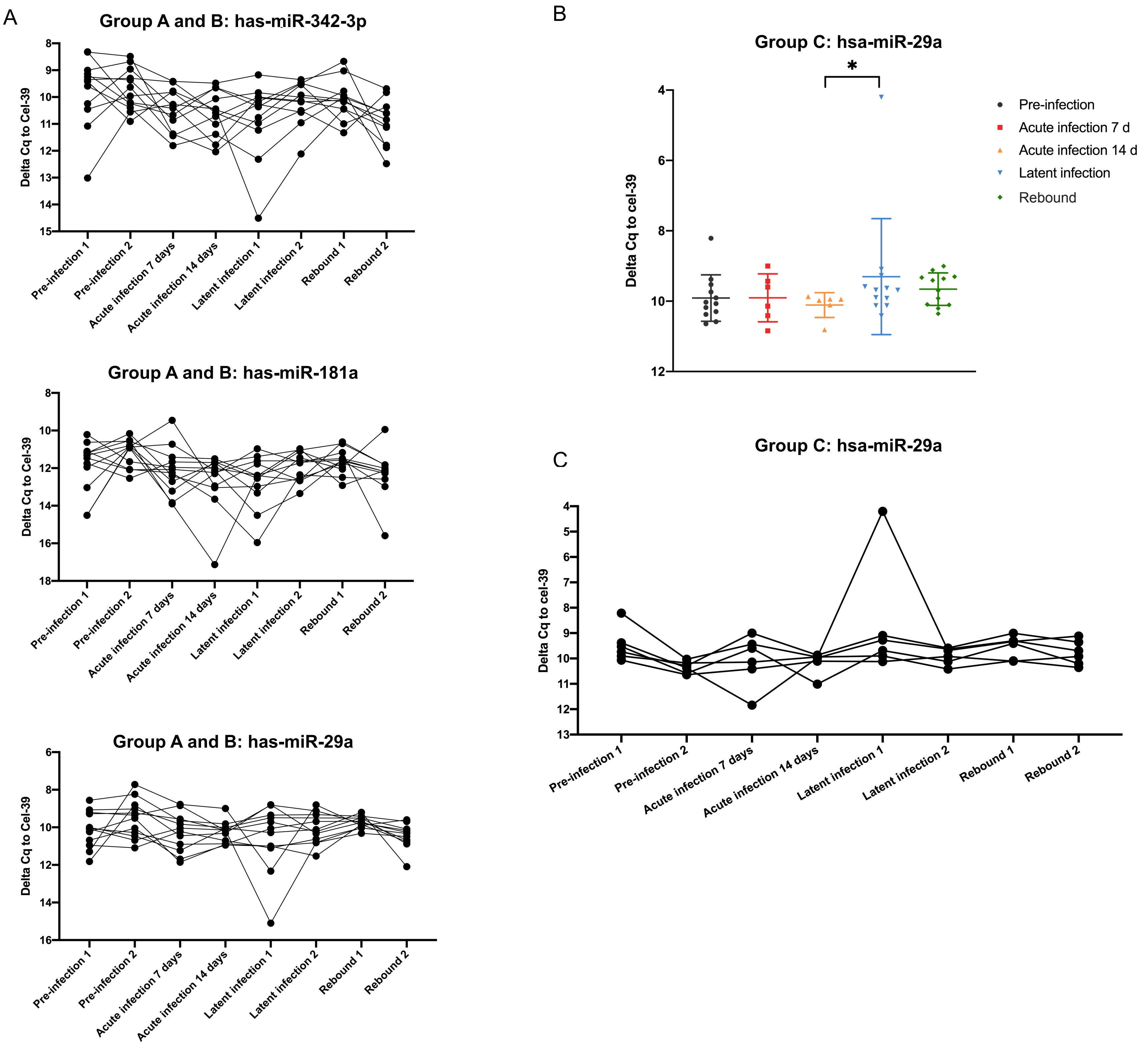
A
Group A and B: has-miR-342-3p
Delta Cq to Cel-39
Pre-infection 1
Pre-infection 2
Acute infection 7 days
Acute infection 14 days
Latent infection 1
Latent infection 2
Rebound 1
Rebound 2
Group A and B: has-miR-181a
Delta Cq to Cel-39
Group A and B: has-miR-29a
Delta Cq to Cel-39
B
Group C: hsa-miR-29a
*
Delta Cq to cel-39
Pre-infection
Acute infection 7 d
Acute infection 14 d
Latent infection
Rebound
C
Group C: hsa-miR-29a
Delta Cq to cel-39
Pre-infection 1
Pre-infection 2
Acute infection 7 days
Acute infection 14 days
Latent infection 1
Latent infection 2
Rebound 1
Rebound 2

Figure S6

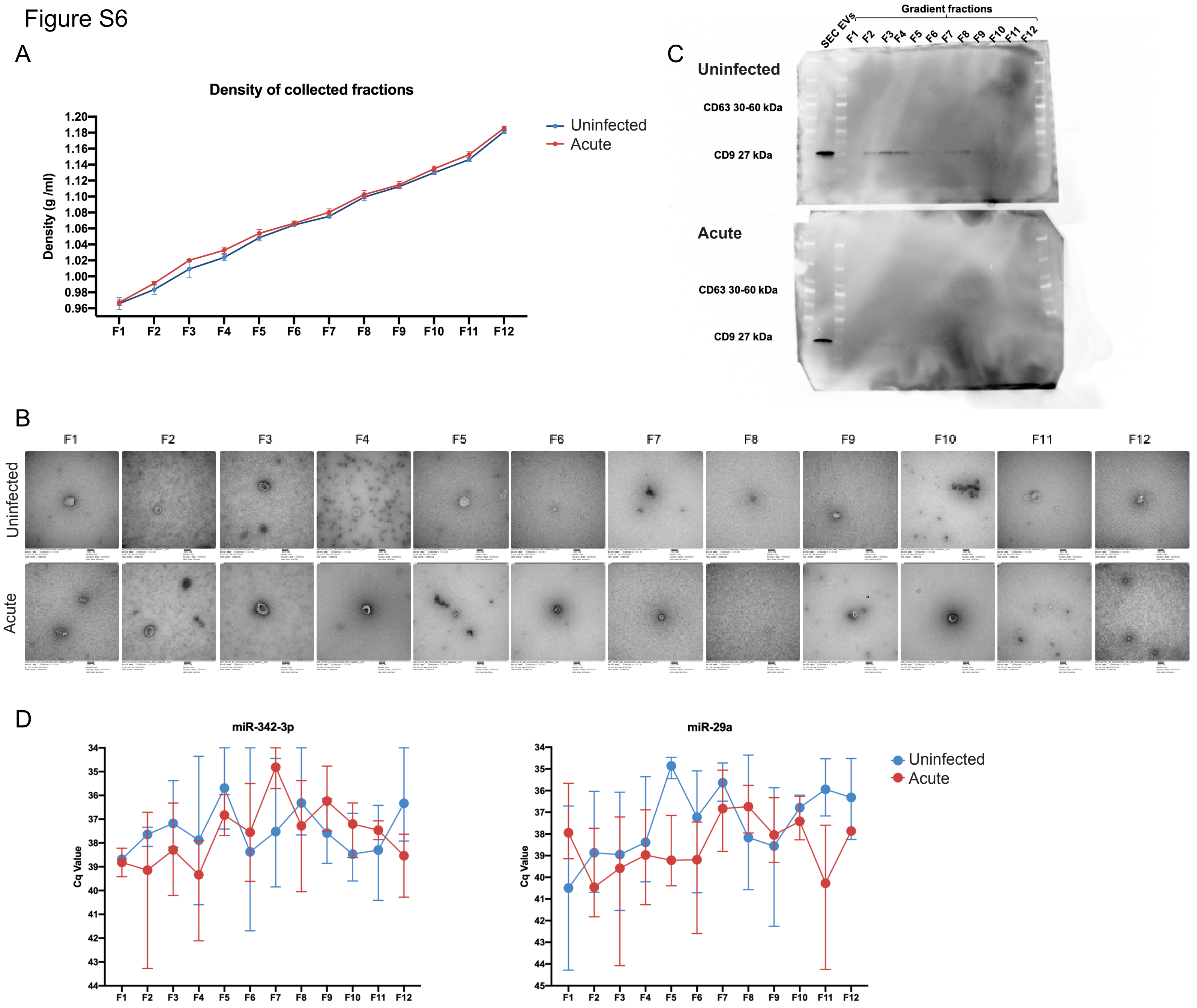
A
Density of collected fractions
Density (g /ml)
1.20
1.18
1.16
1.14
1.12
1.10
1.08
1.06
1.04
1.02
1.00
0.98
0.96
F1 F2 F3 F4 F5 F6 F7 F8 F9 F10 F11 F12
Uninfected
Acute
C
Gradient fractions
SEC EVs F1 F2 F3 F4 F5 F6 F7 F8 F9 F10 F11 F12
Uninfected
CD63 30-60 kDa
CD9 27 kDa
Acute
CD63 30-60 kDa
CD9 27 kDa
B
F1 F2 F3 F4 F5 F6 F7 F8 F9 F10 F11 F12
Uninfected
Acute
D
miR-342-3p
Cq Value
F1 F2 F3 F4 F5 F6 F7 F8 F9 F10 F11 F12
miR-29a
Cq Value
F1 F2 F3 F4 F5 F6 F7 F8 F9 F10 F11 F12
Uninfected
Acute
